# Src/FAK complex phosphorylates cardiac myosin binding protein c (cMyBP-C) in vitro and in vivo

**DOI:** 10.1101/2020.01.28.923870

**Authors:** Ling Wang, Yixin Jin, Jian Wang, Yang Liu, Rui Wang, Shenyuan L. Zhang, Mariappan Muthuchamy, Carl W. Tong, Xu Peng

**Author notes:** Corresponding author: Xu Peng, M.D., Ph.D., Carl W. Tong, M.D., Ph.D.

## Abstract

Cardiac myosin binding protein C (cMyBP-C) is a phosphorylation-dependent force regulator and plays an important role in controlling myosin and actin dynamic interaction. Point-mutations of cMyBP-C that interfere with cMyBP-C threonine/serine phosphorylation resulted in hypertrophic cardiomyopathy and cardiac failure. However, it remains largely unknown how cMyBP-C tyrosine phosphorylation is regulated during cardiac hypertrophy and heart failure. Integrins are receptors of extracellular matrix and are the sensors of cardiac mechanical stretch. Focal adhesion kinase (FAK) plays an essential role in integrin-initiated signal transduction and regulates multiple cellular functions in various types of cells including cardiomyocytes. To identify the regulatory mechanism of cMyBP-C tyrosine phosphorylation during cardiac hypertrophy, we examined the effect of FAK on phosphorylation of cMyBP-C. Immunoprecipitation analysis showed that FAK and cMyBP-C are associated within the intact mouse heart. Results from our mutagenesis experiments demonstrated that the FAK kinase domain was required for FAK to associate with cMyBP-C. Our data also documented that the FAK Y397 site is required for FAK and cMyBP-C association. Importantly, overexpression dominant active Src Y527F with FAK significantly enhanced cMyBP-C phosphorylation. Interestingly, overexpression of cMyBP-C inhibited FAK phosphorylation. Taken together, cMyBP-C is one of effectors of Src/FAK complex in cardiomyocyte.

## Introduction

Cardiac contraction and relaxation rely on actin and myosin cross-bridge cycling, and deregulated cross-bridge cycling impairs cardiac pump functions (1, 2). Cardiac myosin binding protein C3 (cMyBP-C) is a myosin-associated protein and plays an important role in regulating cardiac contraction and relaxation through affecting kinetic interactions between myosin and actin (3). Dephosphorylated cMyBP-C prefer binds to myosin and inhibits the interaction between myosin and actin (4–7). Consistently, the overall phosphorylation levels of cMyBP-C in the heart are decreased in heart failure and cardiomyopathy patients (8–10). In mammalian cells, phosphorylation usually occurs on serine, threonine and tyrosine residues. The importance of serine/threonine phosphorylation of cMyBP-C has been well documented and more than five serine/threonine phosphorylation sites have been identified (11). Recently, newly available information indicates that tyrosine phosphorylation of cMyBP-C may also be involved in cardiomyopathy formation. Mass spectrometry (MS) analysis documented that Tyr79 of cMyBP-C can be phosphorylated (12). Moreover, the Tyr237Ser mutant was reported in hypertrophic cardiomyopathy patients (13). Unlike the important effect of serine/ threonine phosphorylation of cMyBP-C on cardiac function, it remains unclear the physiological functions of cMyBP-C tyrosine phosphorylation during cardiomyopathy formation.

Focal adhesion kinase (FAK) is a non-receptor tyrosine kinase and is critical mediator in integrin-initiated signal transduction (14, 15). FAK contains a centrally located catalytic tyrosine kinase domain and large non-catalytic N- and C-terminal domains (14, 16, 17). Once FAK get activated, the FAK FERM domain, which is located at the N-terminus of FAK, moves away from the kinase domain and induces FAK to switch from an inactive state to an active state (18–20). Tyr397 is the binding site of Src SH2 domain and numerous FAK functions rely on FAK and Src to form a complex (14). In addition, FAK has three proline-rich domains that are important for FAK binding to endophalin A2 and other effector proteins (21).

FAK regulates multiple cellular functions in various types of cells including cardiomyocytes (16, 17). FAK is expressed in both neonatal and adult cardiomyocytes and plays an essential role in regulating embryonic heart development and heart hypertrophy (22–24). FAK is distributed in the multiple subcellular structures of the cardiomyocyte, including the costamere, Z disk and sarcomere (A band) (17). A substantial fraction of FAK is associated with myosin in the sarcomeres under the non-stimulated condition (17). In response to mechanical stress, FAK relocates to the Z-disks and costameres (25). Pulsatile stretch induces rapid FAK activation and FAK phosphorylation is paralleled to the extent and duration of mechanical stimulation, indicating that FAK is involved in mechanical stretch-initiated signal transduction in the cardiomyocytes (26, 27). Once activated, FAK binds to Src through the phosphorylated Tyr397. The FAK/Src complex results in the phosphorylation of Tyr576 and Tyr577 that locate in the FAK kinase domain. Moreover, Src can phosphorylate FAK C-terminal Tyr861 and Tyr925 and create binding sites for other proteins containing SH2 domains. It was also documented that a sustained pressure overload on *in vitro* rat heart induced tyrosine phosphorylation of FAK, as well as enhanced association of FAK with Src and Grb2 (28). We and others have reported that FAK is involved in cardiac hypertrophy formation in response to pressure overload (22, 29, 30). cMyBP-C is an important regulator for myosin and actin cross-bridge cycling and the effect of FAK on cMyBP-C phosphorylation remains enigmatic. In this manuscript, we examined the effect of regulatory mechanisms of FAK on cMyBP-C phosphorylation in cultured cells and pressure overloaded mouse heart.

## Results

### Mechanical stress increased cMyBP-C tyrosine phosphorylation

To determine the effect of pressure overload on cMyBP-C tyrosine phosphorylation, we performed transvers aorta constriction (TAC) or sham surgery on 3-month-old 129/SvJ wild type mice. After 1 hour of TAC surgery, the proximal aorta pressure gradient of the ligation site increased approximately tenfold, compared to the distal ligated aorta (Fig. 1A). Four weeks later, the heart and body weight ratio in the TAC group increased twofold, compared to the sham group (Fig. 1B). To detect the phosphorylation status of cMyBP-C in intact sarcomeres, we isolated myofibers from TAC and sham mice hearts 1 hour later following the surgery. After running an SDS-PAGE, we blotted the membrane with PY100 antibodies that recognize the phosphorylated tyrosine residues. The membrane was then re-probed with cMyBP-C antibodies as a loading control. Our data showed that a pressure overload in the heart significantly increased the level of cMyBP-C tyrosine phosphorylation (around 40%) (Fig. 1C), indicating that cMyBP-C tyrosine phosphorylation may be involved in cardiac hypertrophy in response to pressure overload.

**Fig.1.**
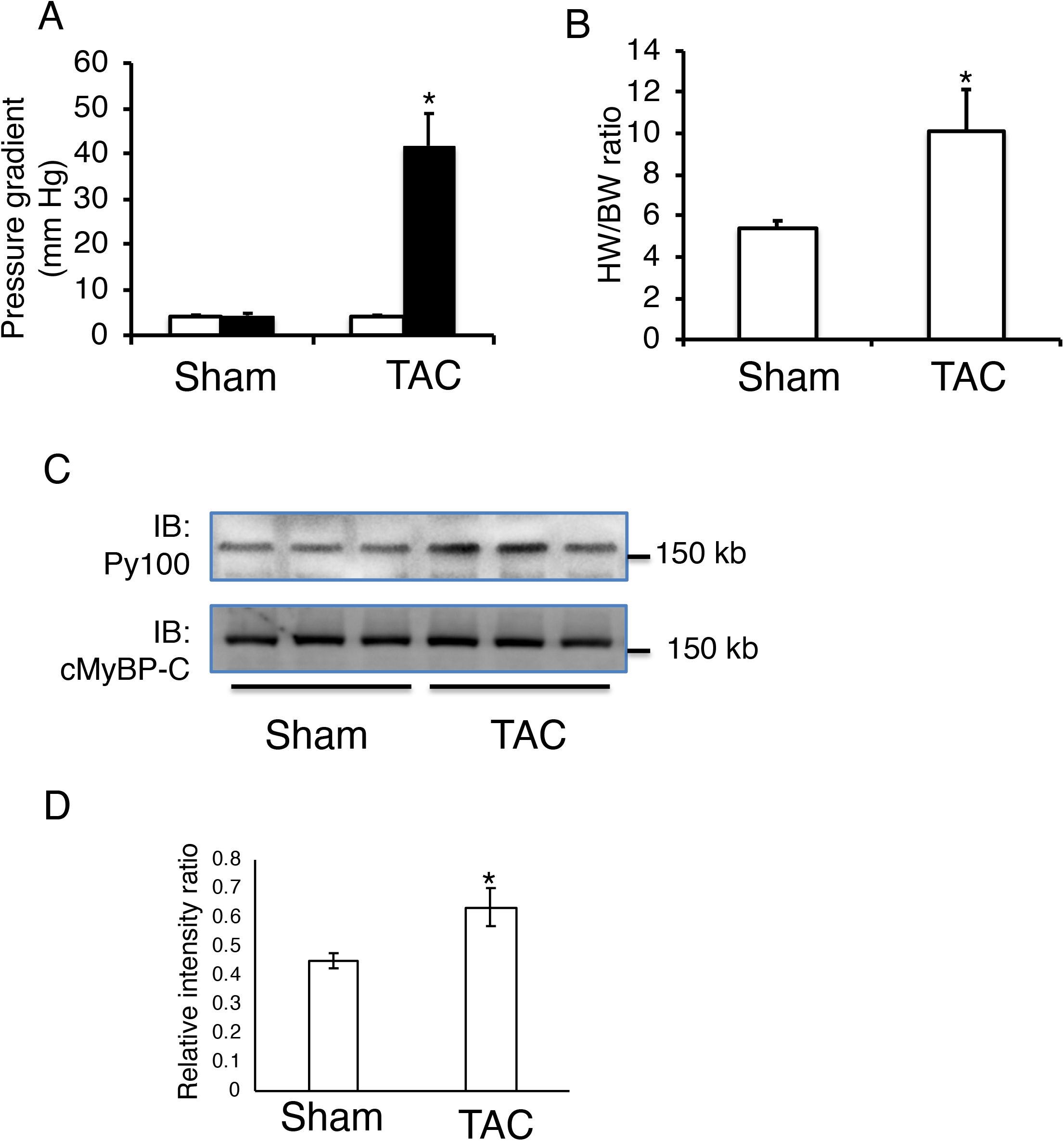
Pressure overload induced cMyBP-C tyrosine phosphorylation in the heart. (A). Measurement of aortic pressure before and 1 hour later after transverse aortic constriction (TAC) or sham surgery. (B). The heart weight to body weight ratio was determined 4 weeks later after TAC or sham surgery. (C). The hearts from TAC and sham mice were harvested and the myofibers were then isolated. The myofibers separated by SDS-PAGE and blotted with Py20 antibodies.

### FAK and cMyBP-C are associated in the intact heart

It is well documented that integrin-mediated signal transduction plays an essential role in pressure overload-induced cardiac hypertrophy (31, 32). FAK is one of the most important tyrosine kinases in integrin-initiated signal transduction, so it is possible that TAC induced cMyBP-C phosphorylation is mediated by FAK. To investigate the potential regulatory mechanisms of cMyBP-C phosphorylation by FAK, we first examined the interaction between FAK and cMyBP-C in the mouse heart. Tissue lysates were prepared from 3-month-old adult mouse cardiac ventricles and then precipitated with FAK antibodies. In parallel, we incubated the tissue lysates with the same amount of control IgG. The Western blot results using cMyBP-C antibody showed that cMyBP-C was associated with FAK, and the normal IgG barely precipitated cMyBP-C (Fig. 2A).

**Fig.2.**
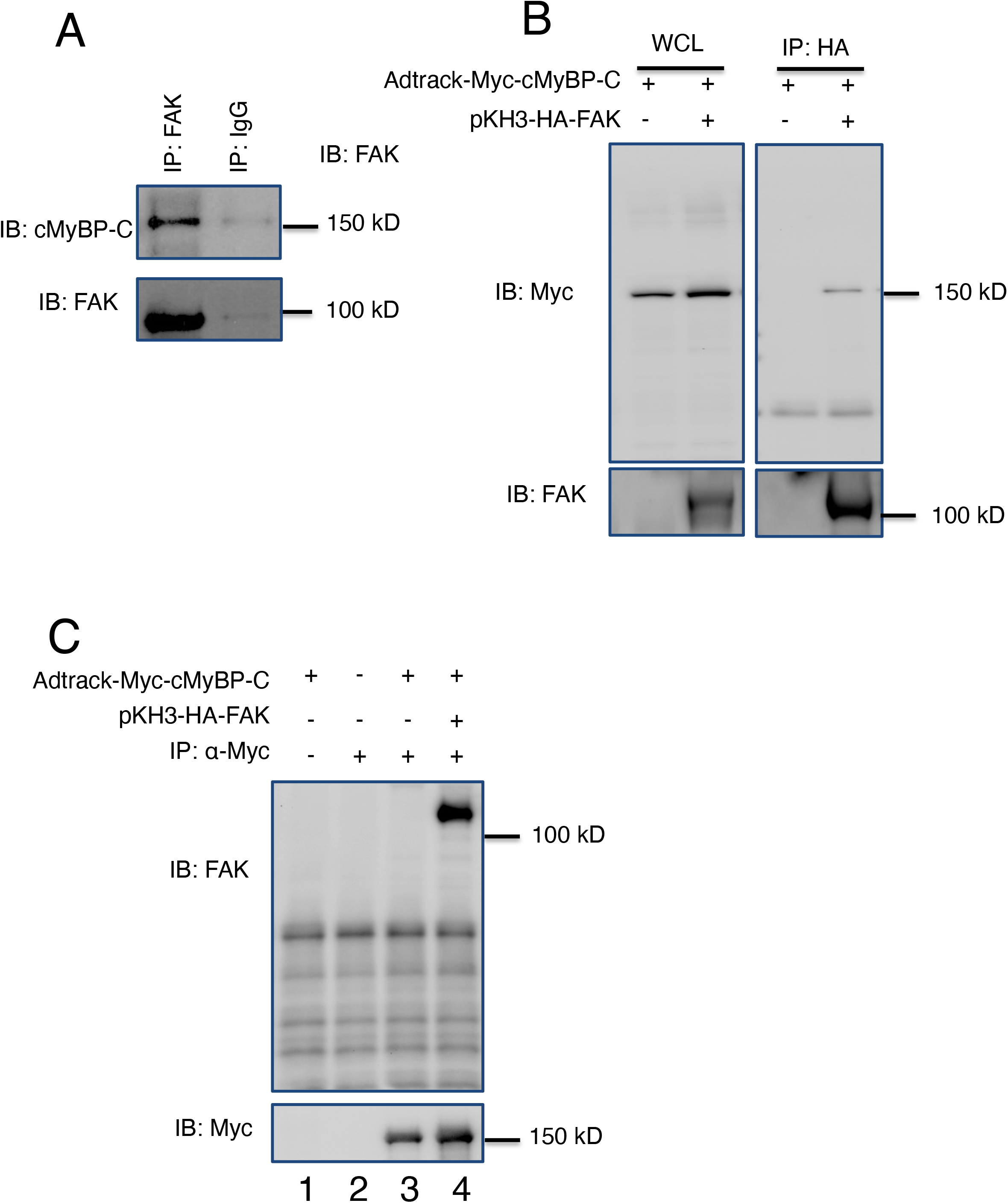
Association of FAK and cMyBP-C. (A). The whole tissue lysates were prepared from 3-month-old wild-type heart. The tissue lysates were incubated with either anti-FAK antibodies or control normal mouse Ig G, as indicated. The precipitates were analyzed by Western blotting with anti-cMyBP-C. (B). Ad 293 cells were co-transfected with Adtrack-Myc-cMyBP-C with/without pKH3-HA-FAK. Whole cell lysates were blotted with anti-Myc and anti-FAK antibodies, as indicated. Ad 293 whole cell lysates were precipated with HA antibodies and analyzed with anti-Myc or anti-FAK antibodies. (C). Ad 293 cell lysates were precipated with anti-Myc antibodies and blotted with anti-FAK and anti-Myc antibodies, as indicated.

To avoid the possibility that the association between FAK and cMyBP-C is through other sarcomere proteins, we overexpressed Myc-tagged cMyBP-C with/without human influenza hemagglutinin (HA)-tagged FAK in Ad 293 cells. The Western blot results showed that the overexpressed cMyBP-C can be detected by the Myc antibody (Fig. 2B, left upper panel), and FAK antibodies recognized the overexpressed FAK and could not detect the endogenous human FAK in Ad 293 cells (Fig. 2B, left lower panel). An immunoprecipitation experiment presented that cMyBP-C was associated with FAK, but not with the pKH3 vector alone (Fig. 2B, right panel). In line with this result, a reciprocal immunoprecipitation experiment showed that FAK was found in anti-Myc immunoprecipitates (Fig. 2C, line 4), but could not be detected in the precipitates by the control normal IgG (Fig. 2C, line 1). Taken together, these results demonstrated that FAK and cMyBP-C are associated in the heart and that this association is not mediated by other sarcomere proteins.

### FAK kinase domain is required for FAK and cMyBP-C interaction

FAK is a non-receptor tyrosine kinase with a central kinase domain flanked by large N- and C-terminal domains (Fig. 3A). To define the cMyBP-C binding domains within FAK, Ad 293 cells were cotransfected with Myc-tagged cMyBP-C with the full-length HA-tagged FAK, HA-FAK N-terminal domain (1-400), HA-FAK-kinase domain (401-664), HA-FAK C-terminal domain (676-1052), and HA-FAK with N-terminus deletion (Δ 1-125). Immunoprecipitations were performed with anti-HA antibodies and were followed by western blotting with anti-cMyBP-C antibodies. Consistent with our previous result (Fig. 2B), cMyBP-C was coprecipitated with the full-length FAK. FAK N-terminus did not associate with cMyBP-C, but the FAK kinase domain was strongly associated with cMyBP-C. Interestingly, we found that the association between FAK mutant (Δ 1-125) and cMyBP-C was increased as compared to the full-length FAK. Similar expression levels of cMyBP-C were verified by blotting of whole cell lysates with anti-cMyBP-C antibodies (Fig. 3B, bottom panel). All these data indicate that the FAK kinase domain is the major binding site for FAK and cMyBP-C association.

**Fig.3.**
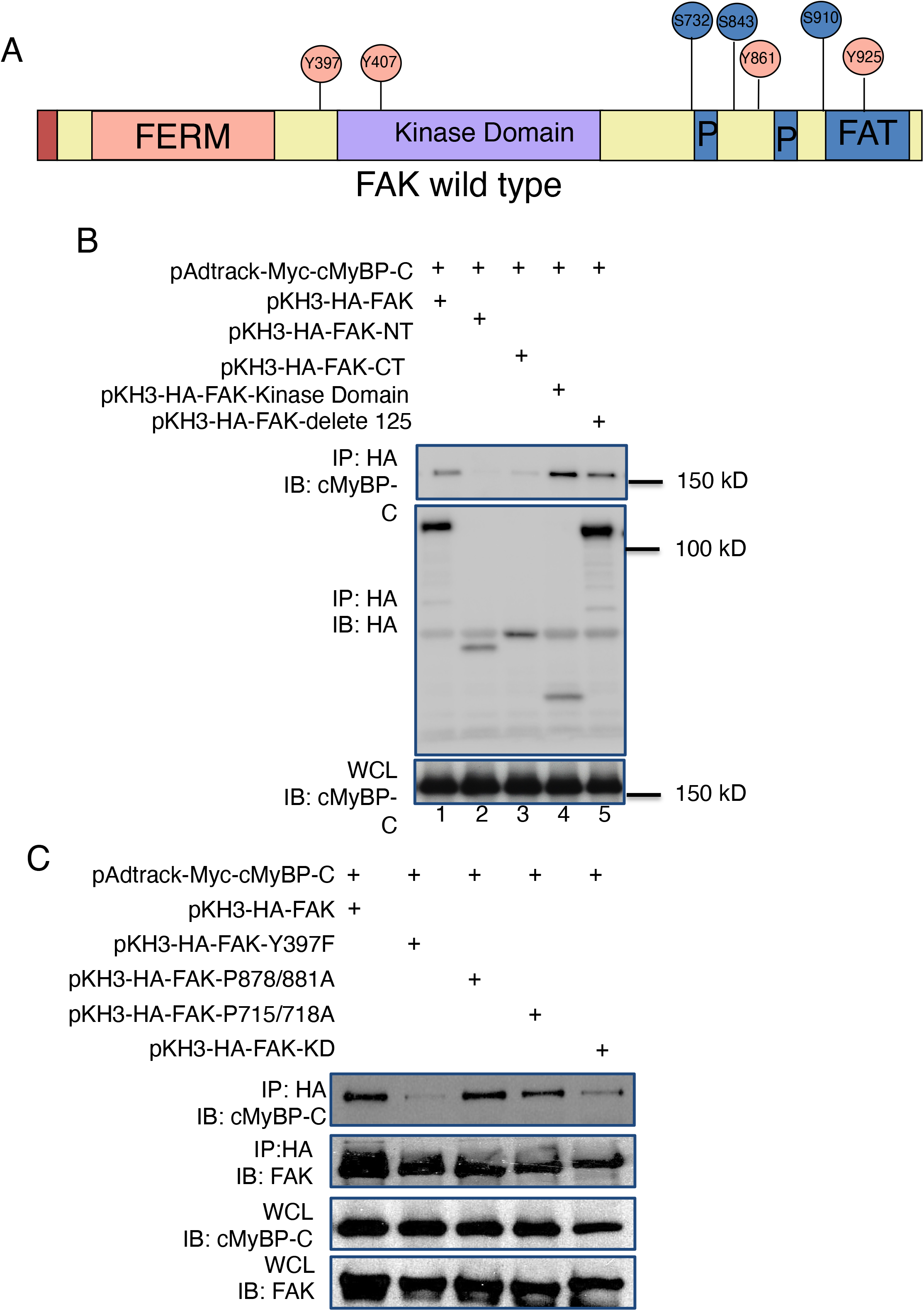
FAK kinase domain and Y397 site are required for the interaction between FAK and cMyBP-C. (A). Schematic diagram of FAK and its fragments used in the association assay in the panel B and C. (B). Ad 293 were transfected with pAdtrack-Myc-cMyBP-C with full length FAK and different FAK fragments as indicated. Cell lysates were precipitated with anti-HA antibodies and were analyzed by blotting with either cMyBP-C or anti-HA antibodies. Aliqous whole cell lysates were analyzed with anti-cMyBP-C antibodies. (D). Ad 293 cells were transfected with pAdtrack-Myc-cMyBP-C with wild type FAK or different FAK point mutations. Cell lysates were precipated with anti-HA antibodies and blotted with either cMyBP-C or FAK antibodies. Aliquots whole cell lysates were blotted with anti-cMyBP-C or FAK antibodies to verify cMyBP-C and FAK expression levels.

As an important signal transduction regulator, FAK binds to different downstream effectors and then regulates various cellular functions. To investigate the potential cMyBP-C binding sites in FAK, Ad 293 cells were cotransfected with cMyBP-C containing different FAK point mutations. Whole cell lysates were precipitated with HA antibodies and followed by western blotting with cMyBP-C antibodies. Our results showed that FAK Tyr397 and kinase activity were required for the associations between cMyBP-C and FAK. However, the FAK Pro878/881Ala mutant and the FAK Pro715/718Ala mutant did not affect the association between FAK and cMyBP-C (Fig. 3C).

### cMyBP-C association with FAK through multiple domains

To determine the regions of cMyBP-C necessary for association with FAK, we created a series of cMyBP-C truncation mutations as shown in Fig. 4A. The Myc tagged cMyBP-C mutants and the full-length of FAK were cotransfected into Ad 293 cells. Three days later, whole cell lysates were precipitated with Myc antibodies and then blotted with FAK antibodies. Our results showed that the full-length cMyBP-C was associated with FAK and that the association of cMyBP-C fragment 4 (F4) with FAK was significantly increased. In addition, we detected that cMyBP-C F5 and F1 fragments were also associated with FAK, indicating that cMyBP-C could associate with FAK through multiple domains (Fig. 4B).

**Fig. 4.**
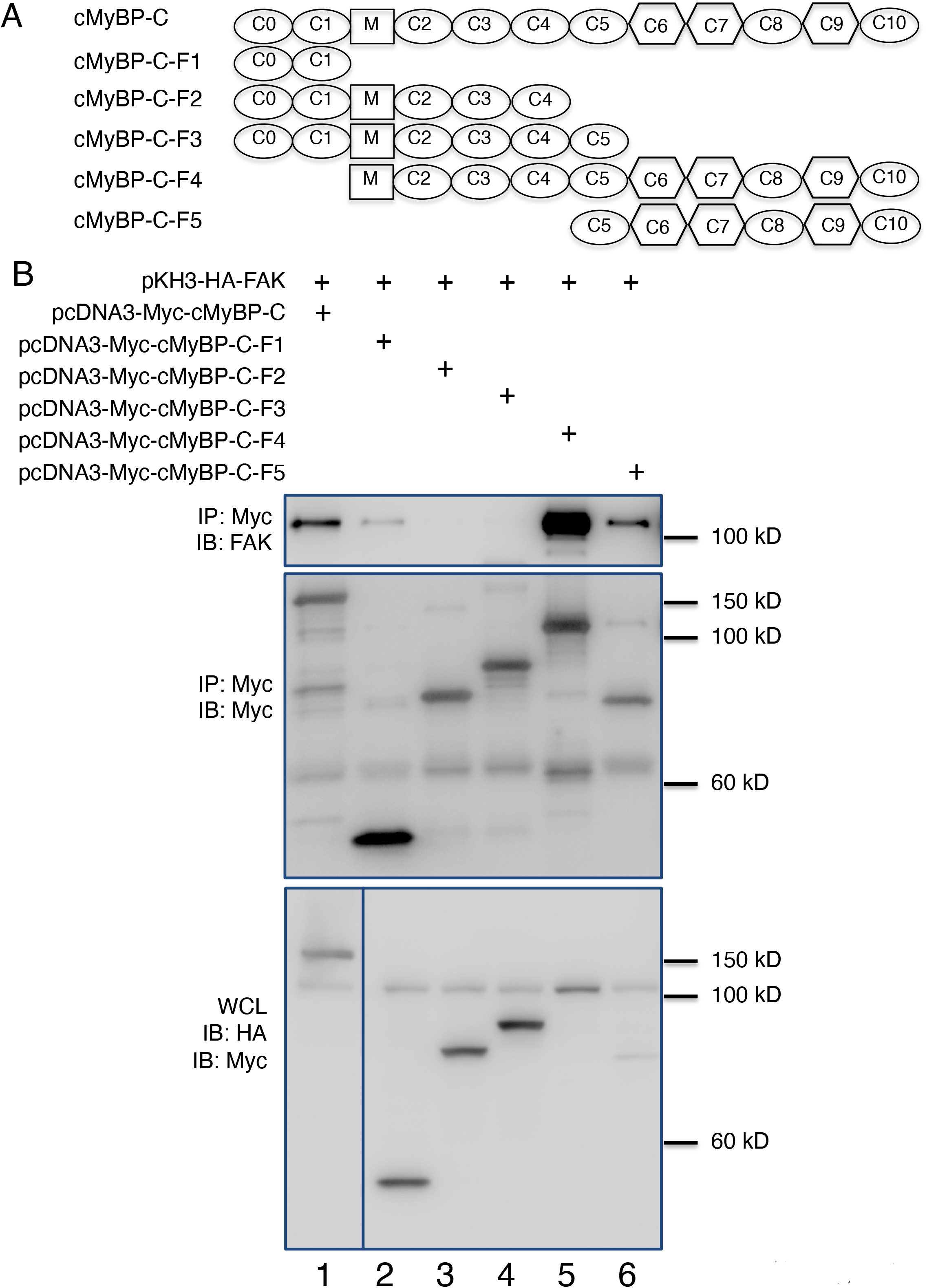
Analysis of cMyBP-C association with FAK. (A). Schematic diagram of cMyBP-C and its fragments used in the association analysis in the panel B. (B). Ad 293 cells were transfected pKH3-HA-FAK with full length cMyBP-C or different cMyBP-C fragments. Immunoprecipatation experiments were performed with anti-Myc antibodies and analyzed with either FAK or Myc antibodies, as indicated. Aliquots whole cell lysates was blotted with anti-HA and Myc antibodies.

### FAK-mediated Src phosphorylation on cMyBP-C

It was reported that Src and FAK can form a complex and then phosphorylate their downstream effector proteins. To investigate the potential physiological significance of association between FAK and cMyBP-C, we examined the effect of Src, FAK and Src plus FAK on cMyBP-C tyrosine phosphorylation. Ad 293 cells were cotransfected cMyBP-C with FAK, V-Src, Src Tyr537Phe, Src plus FAK, and Src Tyr527Phe and FAK. cMyBP-C and FAK expression was confirmed by western blotting using Myc and FAK antibodies, respectively. PY 100 blotting showed that neither v-Src and FAK alone nor v-Src plus FAK could phosphorylate cMyBP-C. The Src Tyr527Phe mutant increased a 150 kD protein phosphorylation. Interestingly, FAK plus Src Tyr527Phe significantly enhanced a 150 kD protein phosphorylation, indicating that FAK and dominant active Src can phosphorylate cMyBP-C (Fig. 5A). To further confirm these results, we performed immunoprecipitation experiments. FAK, v-Src, Src Tyr527Phe with FAK were cotransfected with cMyBP-C into Ad 293 cells. Whole cell lysates were precipitated with Myc antibodies. Western blotting with Myc antibodies showed that the same amount cMyBP-C were precipitated. PY100 blotting showed that the FAK plus Src Tyr527Phe mutant resulted in cMyBP-C phosphorylation (Fig. 5B). Taken together, FAK and dominant active Src formed a complex to induce cMyBP-C phosphorylation.

**Fig.5.**
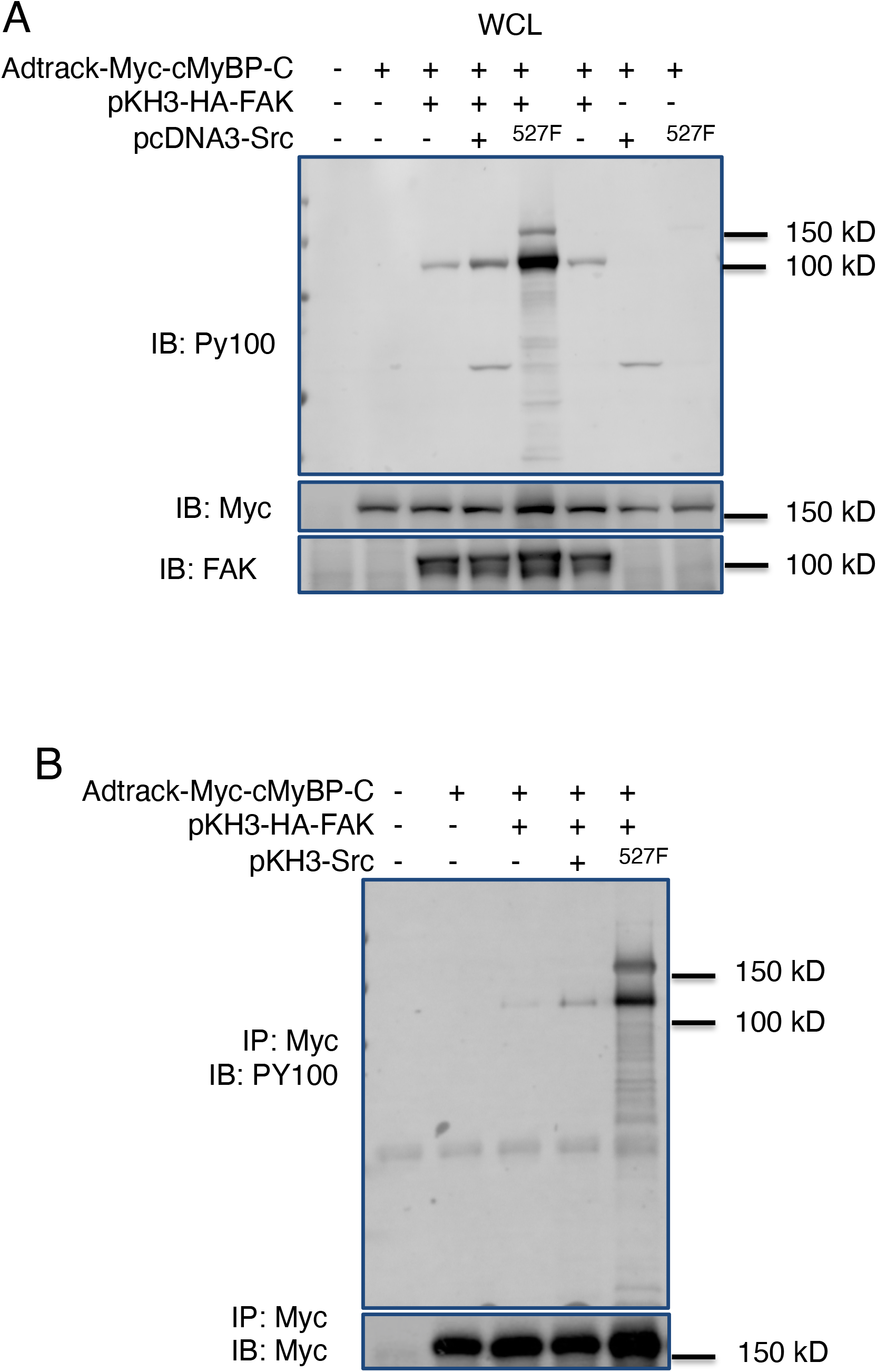
Tyrosine phosphorylation of cMyBP-C by FAK and Src complex. (A). Ad 293 cells were co-transfected with pAdtract-Myc-cMyBP-C with/without pKH3-HA-FAK and pCDNA3-Src or Src 527F mutant. Whole cell lystates were analyzed by blotting anti-PY 100, anti-Myc and anti-FAK antibodies. (B). Immunoprecipatation experiments were performed by using anti-Myc antibodies and analyzed by blotting with anti-PY100 or anti-Myc antibodies, respectively.

### FAK kinase activity is required for cMyBP-C phosphorylation

Previous data showed that FAK and Src Tyr527Phe forming a complex and inducing cMyBP-C phosphorylation. In addition, FAK kinase activity is required for FAK interacting with cMyBP-C. To determine the importance of FAK kinase activity for cMyBP-C phosphorylation, we overexpressed cMyBP-C, Src Tyr527Phe with wild type FAK or FAK kinase dead mutant. In line with previous data, overexpression of wild type FAK or FAK kinase dead mutant alone cannot phosphorylate cMyBP-C. Src Tyr527Phe can weakly phosphorylate cMyBP-C. Overexpression of FAK kinase dead protein and Src Tyr527Phe induced cMyBP-C phosphorylation. Moreover, the phosphorylation level of cMyBP-C was significantly increased when wild type FAK and Src Tyr527Phe were coexpressed with cMyBP-C (Fig. 6A).

**Fig. 6.**
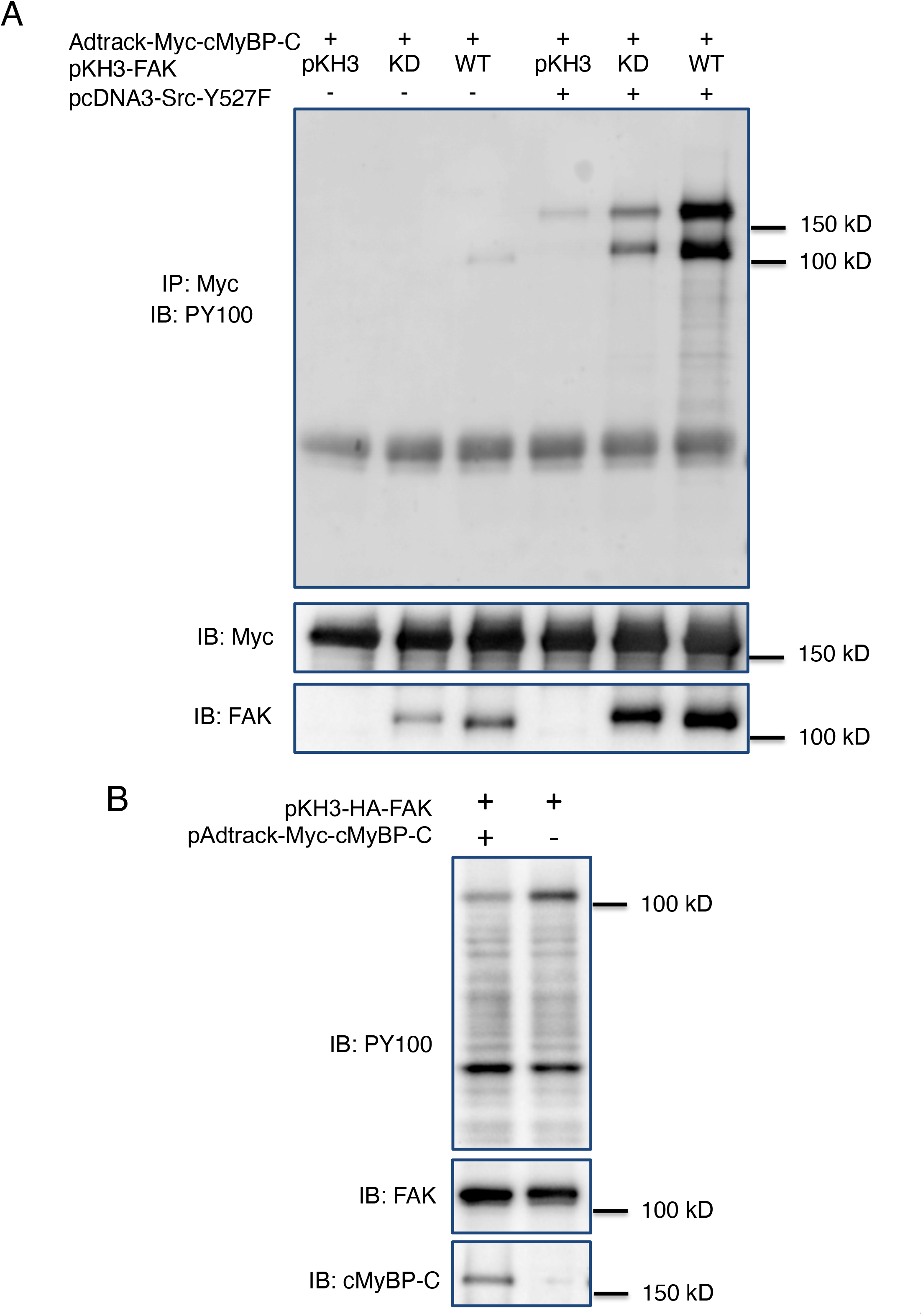
FAK kinase activity is required for cMyBP-C phosphorylation. (A). Ad 293 cells were co-transfected with pAdtrack-Myc-cMyBP-C, pcDNA3-Src-Y525F and pKH3-HA-FAK or FAK kinase dead mutant. Whole cell lysates were immunoprecipatated with anti-Myc antibodies and blotted with anti-PY 100, anti-Myc and anti-FAK antibodies, respectively. (B). Ad 293 cells were co-transfected pKH3-HA-FAK with or without pAdtract-Myc-cMyBP-C. Cell lysates were precipated with anti-HA antibodies and blotted with PY 100, FAK and cMyBP-C antibodies, as indicated.

Because the FAK kinase domain is required for the association between cMyBP-C and FAK, we next examined the effect of cMyBP-C on FAK activity. FAK or the pKH3 vector alone was cotransfected with cMyBP-C and followed by western blot using PY100 antibodies. We found that overexpression of cMyBP-C, instead of the pKH3 vector alone, decreased a 120 kD fragments phosphorylation, indicating that the overexpression of cMyBP-C inhibited FAK phosphorylation (Fig. 6B).

### FAK and Src phosphorylated cMyBP-C *in vitro*

To further determine the role of FAK and Src on cMyBP-C phosphorylation, we overexpressed FAK, Src and FAK plus Src in Ad 293 cells. Immunoprecipitation experiments were then performed using anti-HA antibodies. The purified cMyBP-C was incubated with FAK, Src or the FAK plus Src complex for two hours and then separated by SDS-PAGE. Western blotting results showed that both FAK and Src can phosphorylate cMyBP-C, but the FAK and Src complex did not further enhance cMyBP-C phosphorylation in vitro (Fig. 7).

**Fig. 7.**
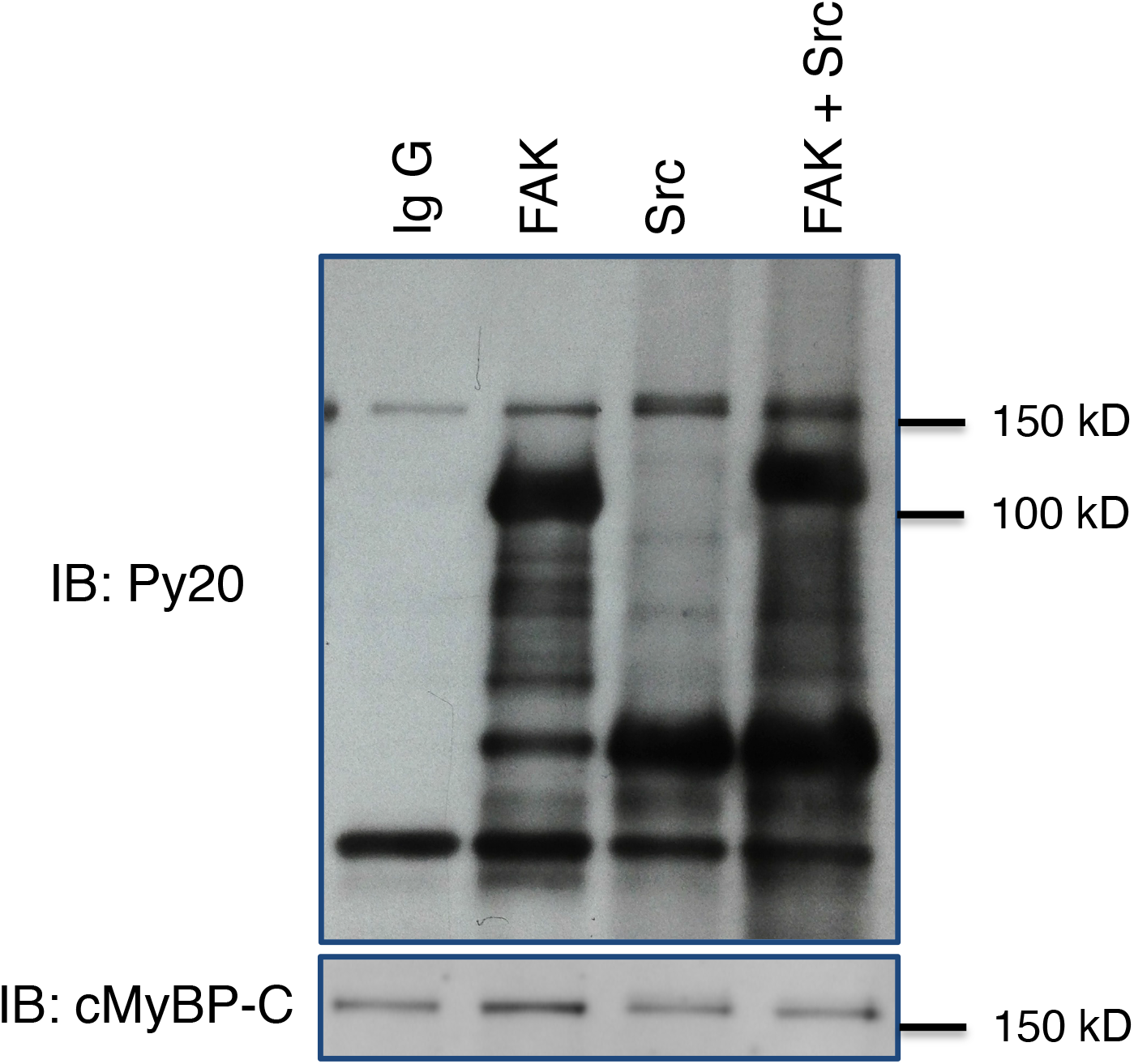
cMyBP-C phosphorylation by FAK and Src. Ad 293 cells were transfected with pKH3-HA-FAK, pKH3-v-Src, or pKH3-HA-FAK with pKH3-HA-v-Src. FAK, Src or FAK + Src were precipitated with anti-HA antibodies and incubated with recombinant cMyBP-C. cMyBP-C phosphorylation was then analyzed by blotting with anti-PY 100 antibodies. The membrane was reprobed with cMyBP-C antibodies to serve as a loading control.

## Discussion

Sarcomere is the contraction unit of cardiomyocyte and its contraction and relaxation functions depend on the dynamic interactions between myosin and actin, which is regulated by cMyBP-C (33). Previous studies have documented that protein kinase A (PKA)-mediated cMyBP-C serine phosphorylation is essential for sarcomere contraction and relaxation (34). However, the regulatory mechanisms of cMyBP-C tyrosine phosphorylation in response to mechanical stretch in the heart remain unclear. In the current manuscript, we have demonstrated that pressure overload enhances cMyBP-C tyrosine phosphorylation in the mice hearts. In addition, we found that the Src/FAK complex phosphorylates cMyBP-C and that the FAK kinase domain is needed for FAK and cMyBP-C association. Moreover, FAK kinase activity is required for the association between FAK and cMyBP-C. We also found that cMyBP-C has multiple domains that bind to FAK and the association between FAK and cMyBP-C inhibited FAK phosphorylation.

Integrin-FAK mediated signal transduction plays an important role in cardiac development and cardiac hypertrophy (22, 23). Previously, we have reported that the inactivation of FAK in cardiomyocytes enhanced TAC-induced cardiac hypertrophy (22). FAK Tyr397 locates at the junction of The FAK N-terminus and the kinase domain and is the binding site for Src and PI3K, etc. (16). Our results showed that the FAK Tyr397Phe mutant disrupted the association between FAK and cMyBP-C. It is possible that cMyBP-C competes with Src and directly binds to FAK Tyr397. This competition may interfere with the association between FAK and Src and decrease FAK phosphorylation. In line with this result, we found that overexpression of cMyBP-C inhibits FAK phosphorylation. Another possibility is that cMyBP-C does not bind to FAK Y397 directly but can only bind to the active FAK kinase domain. In the inactivated stage, the FAK kinase domain was covered by FAK N-terminus. The FAK Tyr397Phe mutant causes FAK to lose its interaction with Src, which resulted in its inability to be fully phosphorylated and expose to the centrally located FAK kinase domain. In this case, cMyBP-C cannot access the FAK kinase domain, which then inhibits FAK and cMyBP-C association.

The cMyBP-C-F4 fragment strongly associates with FAK and its association is even stronger than that of full-length cMyBP-C, suggesting that the C0-C1 fragment inhibits cMyBP-C form association with FAK. We also noted that the association between C5-C10 is much weaker than that of the M-C10 fragment, indicating that M-C4 is required for FAK and cMyBP-C association. Intriguingly, we found C0-C4 fragments do not associate with FAK. A possible explanation is that the C0-C1 fragment inhibits the association between FAK and cMyBP-C.

In response to shear stretch stimulation, FAK relocates to the costermere and Z-disks in cardiomyocyte (35). The Franchini group reported that Myosin binds to the FAK N-terminus and inhibits FAK phosphorylation (36). In another word, the association of Myosin and FAK inhibits FAK activity. Our data shows that cMyBP-C binds to the FAK kinase domain and the FAK/Src complex can then phosphorylate cMyBP-C. Based on this available information, we speculate that pressure overload stimulates FAK to move to the costamere and to form a complex with Src. The Src/FAK complex causes FAK N-terminus move away from its kinase domain. Subsequently, the exposed kinase domain binds to cMyBP-C and stimulates cMyBP-C phosphorylation. The phosphorylated cMyBP-C may change its structure, accelerate myosin and actin interaction and promote cardiac contraction and relaxation. After FAK is dephosphorylated by phosphatases or inhibited by cMyBP-C directly, FAK disassociates from cMyBP-C and then binds to myosin. The interaction between myosin and FAK enhances FAK N-terminus to interact with its kinase domain and then inhibits FAK phosphorylation.

In conclusion, we have demonstrated that FAK can associate with cMyBP-C and that this association is required for the Src/FAK complex to phosphorylate cMyBP-C.

## Acknowledgements

This work was supported by National Institute of Health R01HL145534 to Carl W. Tong and Xu Peng.

## Material and Methods

### The Transverse aorta constriction (TAC) surgery

Transverse aorta constriction (TAC) surgery was performed on 3-month-old wild-type 129/SvJ mice. The mice were anesthetized with 2 % isoflurane with pure oxygen. Bacitracin Ophthalmic Ointment was applied to the eyes and Burprenorphine (0.05-0.1 ug/g) was subcutaneous injected 30 min prior to surgery. A 27-gauge needle was tightened to the aortic arch between the brachiocephalic trunk and left common carotid artery using 6–0 silk suture (CP Medical Portland, OR, USA). Sham-surgery mice underwent an identical procedure except for the aortic ligation. Mice were housed in a pathogen-free facility and all the procedures were approved by the Institutional Animal Care and Use Committee of Texas A&M Health Science Center.

### Echocardiography

Echocardiographic analysis was performed before and after TAC surgery using a VisualSonics Vevo 2100 system (FUJIFILM VisualSonics Inc) to evaluate the pressure gradient of aorta. Mice were anesthetized with 0.5% to 2.5% isoflurane with controlled echo-table temperature at 37°C and real-time monitored ECG. Two-dimensional long and short axis imaging, blood flow Doppler, tissue Doppler, and two-dimensional guided M-mode measurements were obtained using transducer with a frequency of 550 M. The transverse aorta was also visualized with 2-dimensional and color flow imaging. The aortic flow velocity was measured by pulsed wave (PW) Doppler to assess the presence of artery stenosis by TAC through a transducer with a frequency of 250 M. Digital images were analyzed off-line by Vevo2100 software. The pressure gradients were calculated by the Vevo2100 software using the modified Bernoulli equation (pressure gradient = 4* velocity 2). For each study, average value was made from at least three different measurements.

### Immunoprecipitation and Western blotting

Immunoprecipitation and Western blotting experiments were performed as described previously (37, 38). Ad 293 cells were cultured in 6-well plates with DMEM containing 10% fetal bovine serum. Plasmids were transfected into the cells with Lipofectamine 2000 following the manufacture’s instruction. Subconfluent cells or mouse hearts were lysed with modified RIPA buffer (50 mM Tris–HCl, pH 7.5, 150 mM NaCl, 1% NP-40, 1% sodium deoxycholate, 1 mM sodium vanadate, 10 mM sodium pyrophosphate, 10 mM NaF, 1% Triton X-100, 0.5% SDS, 0.1% EDTA, 10 μg/ml leupepetin, 10 μg/ml aprotinin and 1 mM PMSF). Immunoprecipitation was carried out by incubating lysates with various antibodies and protein A beads at 4 °C for overnight. The beads were boiled with loading buffer and resolved by SDS–PAGE. Immunoprecipitation and Western blotting were performed using the following antibodies, rabbit polyclonal FAK antibody (Cell Signaling), HA tag antibody (HA-7, Sigma-Aldrich), Myc tag antibody (A7, Abcam; 4A6 Millipore).

### Generation of plasmids with different cMyBP-C fragments

Myc-tagged cMyBP-C full length and truncated fragments were generated by PCR using the following primers: gag gat cca atg ccg gag cca ggg aag aaa cc and gtg aat tca ctg agg aac tcg cac ctc cag (cMyBP-C full-length); gag gat cca atg ccg gag cca ggg aag aaa cc and gtg aat tca gac agt gag gtt gaa gtt aca gc (cMyBP-C-F1); gag gat cca atg ccg gag cca ggg aag aaa cc and gtg aat tca gtc aat ctt gac ctc cat gaa gtg (cMyBP-C-F2); gag gat cca atg ccg gag cca ggg aag aaa cc and gtg aat tca tgg gac atc gat gac ctt gac tg (cMyBP-C-F3); gag gat cct atg cat gag gcc att ggt tct gga gac and gtg aat tca ctg agg aac tcg cac ctc cag (cMyBP-C-F4); gag gat cca atg ttt gtg cct agg cag gaa cct ccc and gtg aat tca ctg agg aac tcg cac ctc cag (cMyBP-C-F5). PCR products were digested with BamH I and EcoR I and then inserted into pCDNA3-Myc vectors.

### Myofibril and cytosolic fraction Preparation

Hearts were homogenized using a motor-driven homogenizer in 2 ml of ice-cold K60 buffer (60 mM KCL, 20 mM MOPS at pH7.4 and 2 mM MgCl2) with protease inhibitor (Sigma P8340, 1:100 dilution), 10 uM phosphatase inhibitor (Sigma) and 1mM EDTA. The homogenate was then centrifuged at 1000g for 5 minutes at 4°C. After incubation the supernatant with 1% Triton-X 100 for 30 minutes on ice, the whole tissue lysates were centrifuged again at 1000g for 10 minutes at 4°C. The supernatant was transferred to a clean Eppendorf tube and kept as cytosolic fraction. The pellet was washed twice with K60 buffer containing protease inhibitor and re-suspended in K60 with 1ug/ul of BSA. The concentration of protein was evaluated with Piece® BCA Protein Assay Kit.

### Statistical analysis

Data are presented as mean ± standard deviation. Means were compared by analysis of variance between groups (ANOVA). P≤0.05 was considered statistically significant.

